# *In vivo* tracking of *ex vivo* generated ^89^Zr-oxine labeled plasma cells by PET in a non-human primate model

**DOI:** 10.1101/2024.05.24.595782

**Authors:** David J Young, Abigail J Edwards, Kevin G Quiroz Caceda, Ella Liberzon, Johana Barrientos, Sogun Hong, Jacob Turner, Peter L. Choyke, Sean Arlauckas, Adam S Lazorchak, Richard A Morgan, Noriko Sato, Cynthia E Dunbar

## Abstract

B cells are an attractive platform for engineering to produce protein-based biologics absent in genetic disorders, and potentially for the treatment of metabolic diseases and cancer. As part of pre-clinical development of B cell medicines, we demonstrate a method to collect, *ex vivo* expand, differentiate, radioactively label, and track adoptively transferred non-human primate (NHP) B cells. These cells underwent 10- to 15-fold expansion, initiated IgG class switching, and differentiated into antibody secreting cells. Zirconium-89-oxine labeled cells were infused into autologous donors without any preconditioning and tracked by PET/CT imaging. Within 24 hours of infusion, 20% of the initial dose homed to the bone marrow and spleen and distributed stably and equally between the two. Interestingly, approximately half of the dose homed to the liver. Image analysis of the bone marrow demonstrated inhomogeneous distribution of the cells. The subjects experienced no clinically significant side effects or laboratory abnormalities. A second infusion of B cells into one of the subjects resulted in an almost identical distribution of cells, suggesting a non-limiting engraftment niche and feasibility of repeated infusions. This work supports the NHP as a valuable model to assess the potential of B cell medicines as potential treatment for human diseases.

## Introduction

Enzyme replacement therapies (ERT) and antibody therapies have revolutionized outcomes in conditions such as hemophilia, von Willebrand disease, lysosomal storage disorders, metabolic disorders, immunodeficiencies, and cancer. ^1–7^ Yet these therapies suffer from the inconvenience of frequent parenteral administrations, infusion reactions, disease breakthrough, immune rejection, and expense. Gene therapy offers a stable alternative to ERT, eliminating the need for life-long treatments, however, these approaches are also subject to concerns about delivery, limitations of gene editing, loss of edited cells, and immune intolerance.^8–10^ Potentially curative hematopoietic stem cell transplantation (HCT) with allogeneic or genetically modified autologous hematopoietic stem cells (HSC) requires niche clearance via conditioning modalities such as chemotherapy, with associated acute toxicities and long-term complications.^11^ Delivery of a therapeutic protein via a long-lasting engraftable cellular factory not requiring conditioning or human leukocyte antigen (HLA)-matching is a desirable alternative.

Plasma cells are terminally differentiated antibody-secreting progeny of B cells capable of generating tens of thousands of immunoglobulin molecules each second and producing gram-scale quantities of protein on a continuous basis.^12^ Quiescent, long-lived plasma cells arise from terminally differentiated B cells in lymphoid tissues and home to specialized niches such as bone marrow, persisting for years to decades.^13,14^ Following ABO blood type incompatible allogeneic HSC transplantation, donor-specific anti-red blood cell antibodies can persist for years, with documentation of responsible residual recipient plasma cells persisting even in the absence of detectable recipient B cells.^15^ Recipient-derived humoral immunity to pathogens can also persist long-term following HCT.^16^ This post-HCT persistence demonstrates another desirable quality of plasma cells: immunologic “stealth”. Long-lived plasma cells have low major histocompatibility complex expression, allowing them to evade adaptive immunity and resultant allogeneic rejection, even without preconditioning or immunosuppression.^17^ This suggests the possibility of off-the-shelf cellular therapies without HLA-matching.

B cells are highly abundant and easily collected from peripheral blood without mobilization.^18,19^ Primary human B cells can be many-fold expanded *in vitro* and differentiated to antibody secreting plasmablasts and mature, isotype-switched plasma cells. Challenges to B cell medicine approaches include inefficient transduction, achieving protein production at a level likely to be therapeutic, and manufacturing at scale.^20^ However, recent breakthroughs in B cell editing^14,21–24^ using CRISPR/Cas9 coupled with adeno-associated virus mediated delivery of a homology directed repair template enables the efficient introduction of transgenes into safe-harbor loci (e.g., *CCR5*) or the immunoglobulin locus.^22–25^ Advances have also been seen with lentiviral systems.^21^ These techniques produce stable expression at levels predicted to be clinically significant.^25–27^ Furthermore, xenotransplant models confirm that edited human plasma cells derived from *ex vivo* expanded B cells can be detected for up to a year following infusion and continue to produce human immunoglobulins detectable in the plasma.^14^ As a result of these discoveries, work is underway to engineer B cells to produce clotting factors,^23^ immunotherapeutics,^22^ pathogen-specific antibodies for HIV and RSV,^21,25,26^ and viral inhibitors.^25^

Traditionally, the mouse is the platform of choice for pre-clinical experimentation, including studies to date on the adoptive transfers of engineered B lineage cells, due to ease of manipulation and well-characterized genetic background.^28–30^ Xenotransplantation of human cells into immunodeficient mice permits direct assessment of potential therapeutic products, however these mice cannot be used to assess acute infusional or inflammatory toxicities, nor the immunogenicity of residual editing components. In addition, the trafficking, engraftment localization, and persistence of infused human B cell lineage cells may be altered due to the lack of native niches and fully cross-reactive supportive cytokines, a drawback of the cross-species model. Humanization of mice with plasma cell supportive cytokines such as human IL-6 or autologous hematopoietic cells is necessary to support marrow plasma cell engraftment or trafficking to and persistence in the spleen.^14,31^ Finally, demonstration of manufacturing scalability and safety of delivery is critical prior to establishing a clinical program, which mouse models cannot deliver.^28^ These considerations may be relevant for regulatory review.^32^

Adoptive cell therapy modeling in non-human primates (NHP) such as the rhesus macaque (RM, *M. mulatta*) can address many of these preclinical questions.^28,29^ NHP immune function (innate and adaptive) and the architecture of marrow and secondary lymphoid tissues are highly homologous to humans. In many cases, *ex vivo* expansion procedures developed for human therapies can be directly applied to RM cells, due to cross-reactivity of cytokines and similar cellular properties. NHP models with intact immune systems have been demonstrated to recapitulate many sensitivities and toxicities encountered with various cellular therapies in clinical use, notably CAR-T cells.^33–36^ Finally, some of the practical difficulties NHPs present for routine research use are a benefit in late preclinical development: longevity allows prolonged longitudinal follow-up, large size facilitates modeling of scaling up for cell production, and outbreeding simulates the variability encountered in clinical trials.^37–39^

Previously, our group has developed zirconium-89 (^89^Zr)-oxine as an ex vivo cell labeling agent for tracking cells in vivo by positron emission tomography (PET) imaging in order to study localization of various types of transplanted hematopoietic cells in mouse and RM recipients.^40–42^ These prior studies indicate that ^89^Zr-oxine labeling does not impact biological function of hematopoietic cells, including natural killer, dendritic, cytotoxic lymphoid, and stem cells.^40–42^ Infusion of deferoxamine chelates ^89^Zr released by cell death and enhances renal excretion, thereby inhibiting bone uptake of unbound ^89^Zr. This method allows high sensitivity detection of live labeled cells, on the order of thousands of cells per cubic centimeter, due to the low radioactive signal in the recipient.^40–42^ With a half-life of 3.3 days, ^89^Zr activity is detectable for up to two weeks following administration of labeled cells; however, the need for increasing scan time to compensate for the radioactivity decay limits practical whole-body imaging to approximately one week. PET images can be overlaid on simultaneous CT images for better anatomic localization. In this report, we combine ^89^Zr-oxine in vivo live-cell imaging technology with breakthroughs in B cell manipulation to establish a pre-clinical RM model of B cell medicines, allowing tracking of the efficiency of engraftment, tissue localization, and persistence for up to a week. This work provides a platform to study the pre-clinical effects of B cell medicine delivery, engraftment/biodistribution, and safety of delivered cells.

## Results

### Collection and *ex vivo* expansion of rhesus macaque B cells

Peripheral blood mononuclear cells were collected from healthy adult RM via non-mobilized apheresis (**Figure 1A**). CD20-positive B cells were enriched using immunomagnetic selection, resulting in greater than 90% purity, and cryopreserved as starting material for the studies described herein. RM peripheral blood B cells have been successfully expanded and differentiated *ex vivo* into class switched antibody secreting cells using commercially available B cell expansion media.^43,44^ We optimized these culture conditions for *ex vivo* B cell RM expansion and differentiation and were able to consistently achieve 10- to 15-fold cell expansion over 7 days, with preserved viability (80-90%) starting with cryopreserved peripheral blood B cells from multiple subjects (**Figure 1B**). After 7-8 days of culture, B cell expansion slowed, and viability began to decline to below 80%.

**Figure 1.**
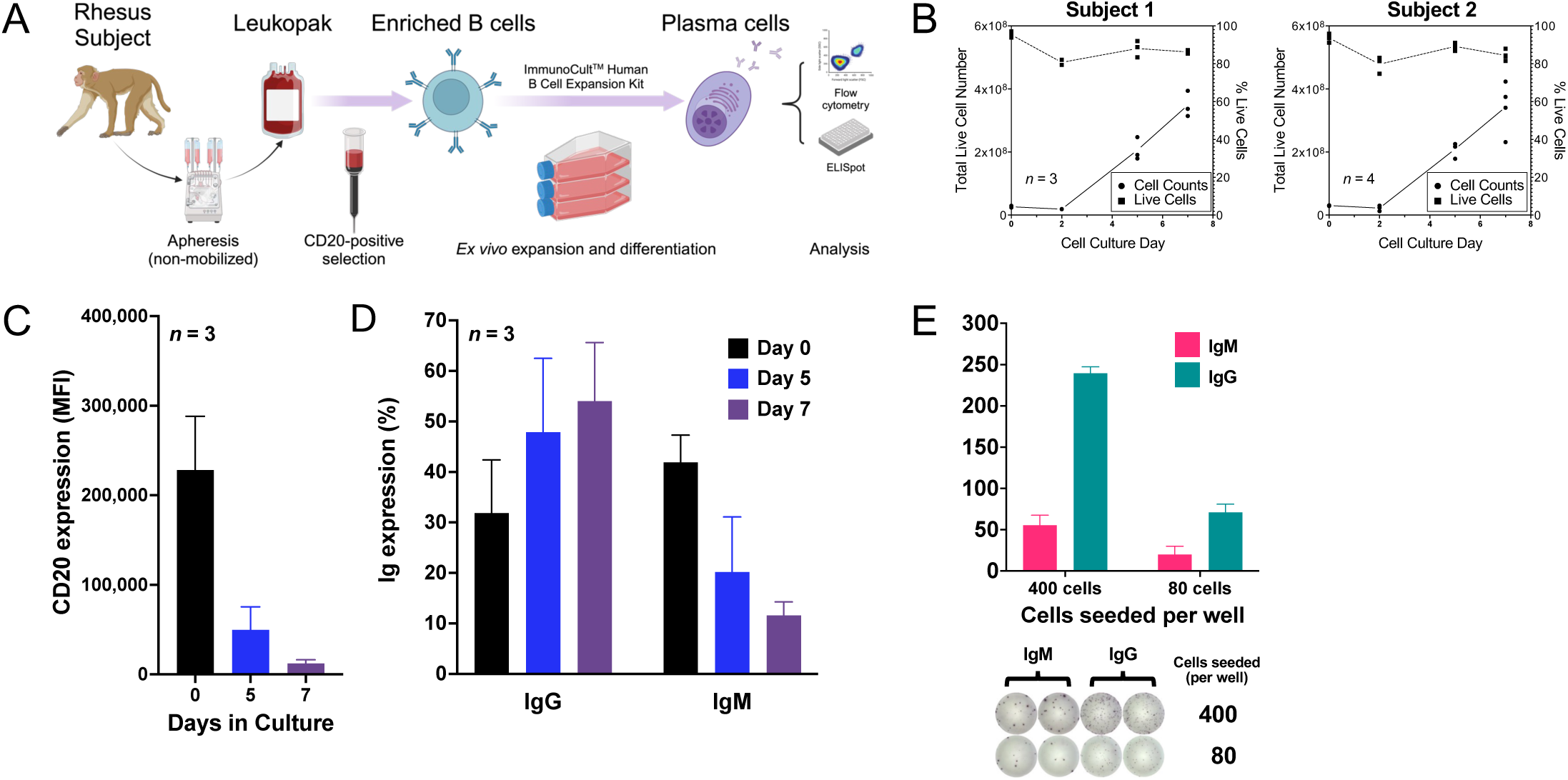
Collection and *ex vivo* expansion of rhesus macaque B cells. A. Schema for the collection, selection, and expansion of RM B cells. B. Expansion and viability of RM B cells from two donors during *ex vivo* culture. Total cell number (primary/left axis) and cell viability as determined by acridine-orange/propidium iodide staining (secondary/right axis) are plotted as a function of time in culture. The plots show the average of at least three separate expansions. C. Flow cytometric analysis of B cells over time in *ex vivo* culture demonstrating gradual loss of CD20 expression. Mean fluorescence intensity (MFI) is plotted as a function of time, averaging across repeat experiments (*n* = 3). D. Flow cytometric analysis demonstrates robust expression of IgM shifting to IgG (class switching) during *ex vivo* expansion of RM B cells (*n* = 3). E. A representative ELISpot analysis with quantification demonstrating class switching from production of IgM to IgG after 7 days in culture. Error bars represent standard deviation of experimental replicates in each graph.

### *Ex vivo* expanded macaque B cells demonstrate normal differentiation, class-switching, and immunoglobulin secretion

Mature B cells express surface CD20, which is downregulated upon differentiation into plasmablasts and plasma cells.^45^ *Ex vivo* cultured RM B cells began to down-regulate the expression of CD20 after 2-3 days and were largely CD20-negative by day 7 (**Figure 1C**). The acquisition of surface CD138 expression associates with plasma cell differentiation and was observed on a fraction of the cultured cells on culture day 7 and was further upregulated by replacing the human B cell expansion supplement with medium containing IL-2, IL-6, IL-10, and IL-15 for 3 days, and then with medium containing IL-6, IL-15 and IFN-α 2b for 2 days (**Supplemental Figure 1**). During the second week of *ex vivo* culture, proliferation ceased, and viability decreased significantly, consistent with terminal differentiation into plasma cells. At culture initiation, more than 50% of the CD20-positive RM B cells expressed IgM (**Figure 1D**). After 7 days of *ex vivo* expansion, the proportion of IgM^+^ cells declined and a corresponding increase in IgG^+^ cells was observed, indicating that Ig class switching was occurring (**Figure 1D**). ELISpot analysis on day 7 confirmed that IgG class-switched antibody secreting cells were present in the culture (**Figure 1E**). The high cell yield, high viability, and presence of differentiated, class-switched antibody secreting cells in the day 7 cultures prompted us to explore the engraftment and biodistribution of these cells when infused back into the autologous subject.

### ^89^Zr-oxine labeling of rhesus macaque *ex vivo* expanded B cells

^89^Zr-oxine is a radioactive tracer for PET that readily permeates cells and transchelates to cellular proteins (**Figure 2A**).^42^ Expanded RM B cells were labeled with ^89^Zr-oxine tracer with approximately 50% efficiency (incorporated activity versus initial activity) without significant loss of viability (<10% decrease in viability compared to baseline prior to labeling; 80-90% viability after labeling). The high viability and yield minimize the influence of radiolabeled dead cells upon subsequent imaging analyses. We expanded RM B cells for 7 days and performed ELISpot analysis before and after ^89^Zr-oxine labeling for each infusion product (**Figure 2B**). No concerning differences were noted between labeled and unlabeled cells regarding immunoglobulin production for any infused cell product.

### Autologous infusion of labeled, *ex vivo* expanded, macaque B cells

We infused ^89^Zr-oxine labeled, expanded autologous B cells into two animals followed by serial PET/CT imaging (**Figure 3A**). The animals received no pre-conditioning to facilitate engraftment. After initiation of deferoxamine infusion, Subject 1 (ZJ34), a 12-year-old female RM, received 10 x 10^6^ cells/kg (7.97 x 10^7^ total) with an activity of 4.81 MBq (60 mBq/cell) and Subject 2 (13U002), a 10-year-old male, received 6 x 10^6^ cells/kg (6.64 x 10^7^ total) with an activity of 2.15 MBq (32 mBq/cell). The animals did not experience any clinically concerning reactions or toxicities from the infusions (**Supplemental Figure 2**), with only mild, transient increases in hepatic enzymes, blood urea nitrogen, creatinine, and temperature shifts typically observed following the prolonged and multiple periods of anesthesia necessary for the study procedures and imaging. All lab values normalized within several days. Following cell infusion, the subjects were imaged via PET/CT at several time points on the day of infusion, and then at approximately 24, 72 and 144 hours after infusion (**Figure 3B**).

We observed rapid initial accumulation of cells within the lungs upon initial PET imaging at 5-15 minutes, decreasing by one hour, and disappearing by 24 hours (**Figure 3B**, Subject 1 and Subject 2, Infusion 1). Beginning at 5-15 minutes, and increasing by 1 hour and then 1 day, cells were detected in both liver and spleen. A faint initial localization to the marrow was observed by one hour, increasing by 1 day. By 24 hours, cell distribution in the marrow, spleen and liver reached a steady state, followed by a modest decline over the 6-day observation period. The liver had the most total activity at all time points starting 1 day after infusion, owing to its large volume, whereas the spleen had the highest signal density and thus cell concentration per volume tissue.

### Reinfusion of autologous, labeled, *ex vivo* expanded non-human primate B cells

Although delivered B lineage cells would be hypothesized to be long-lived, over the course of a subject’s lifetime, it may be necessary to “boost” B cell/plasma cell adoptive therapies via reinfusion of autologous cells, particularly when treating children. We asked whether re-infusion would result in the same distribution and dynamics, or if a potential hurdle would be competition for limited niches, particularly in the bone marrow. A second expansion and differentiation of B cells from Subject 2 was performed, with confirmation of phenotype and immunoglobulin production (**Figure 2B**). The cells were labeled with ^89^Zr-oxine. 70 days following the first infusion, 9.6 x 10^6^ cells/kg (1.07 x 10^8^ total) with an activity of 3.55 MBq (33 mBq/cell) were infused (**Figure 3B**, Subject 2, Infusion 2). The pattern of cell distribution was virtually identical to that of the first infusion, with rapid localization to the lungs, followed by redistribution of the cells to the liver, spleen, and marrow by day +1. Again, no clinically significant changes in vital signs or laboratory values were noted during or following the second infusion (**Supplemental Figure 2**).

### Quantitative analysis of the distribution of *ex vivo* expanded cells following infusion

Analysis of activity distribution on the acquired PET images allows quantitative evaluation of cellular distribution. With correction for decay of ^89^Zr over time (approximately 2 half-lives in this 6-day observation period), it is possible to estimate the cellular density of the infusion product within volumes-of-interest (VOI) down to the sub-organ level. With a voxel size of 2.73 x 2.73 x 3.27 mm (24.4 mm^3^) and a starting specific activity of approximately 25 mBq/cell, cell densities as low as 4,000-10,000 cells/cm^3^ can be detected and measured. Of note, these measurements define the *minimum* labeled cell density present. Cell death and turnover could reduce the observed activity within an organ at a rate greater than radioisotope decay, compounded by high protein secretion (i.e., immunoglobulin) by this specific cell type soon after infusion of labeled cells – given that ^89^Zr-oxine labels intracellular proteins – decreasing the decay-corrected specific activity of each labeled cell. Nonetheless, as the goal of these experiments is to demonstrate trafficking and at least short-term stability of engrafted *ex vivo* expanded cells, these minimum estimated cell densities provide clinically and pharmaceutically relevant information.

During the hour following administration and at one hour, >90% of the infused activity can be accounted for (**Table 1**), with the vast majority (88-93%) found in the lungs, liver, spleen, and bone marrow (**Table 2**, **Figure 4A**). One day following infusion, 78-83% of the initial activity remained, with 90% in the liver, spleen, and marrow, accounting for 72-75% of the initial dose. By 24 hours, the lung activity was approximately 1% of the entire total body activity and did not contribute significantly for the remainder of the observation period. The spleen and marrow contained approximately 11-14% and 8-11% of the initial activity infused (10-15% and 13-17% of the total body activity), respectively. 48-54% of the infused activity (63-65% of the total body activity) was found within the liver at 24 hours. The total organ activity as a fraction of the total body activity remained relatively unchanged for these organs. Accounting for infusion product specific activity and correcting for radioisotope decay, these observations can be expressed as the minimum cell occupancy in each organ system (**Table 1**). Although the cell densities in the various organs were markedly higher in the Subject 2’s second infusion compared with the first infusion, this reflects the higher cell dose administered (**Table 1**). Notably, the relative distribution of cells and their kinetics were almost identical to the first infusion (**Table 2**, and **Figure 4A-B**).

As the PET signal was inhomogeneous, we parsimoniously reconstructed the VOI to include only those anatomic regions with clearly delineated signal (**Supplemental Figure 3**), and restricted bone marrow analysis to the vertebral bodies, pelvis, and proximal long bones (humeri and femurs), given restrictions on scan times (see below), and the known replacement of distal marrow by fat with aging in primates (human and RM), in contrast to virtually 100% lifelong cellular marrow in rodents. We estimated the cell density for each organ system over time (**Figure 4C**, **Table 3**). Most notably, across the observation period, the spleen had a ten- to fifteen-fold greater density of cells than the bone marrow, reflecting the spleen as a large lymphoid organ, compared with the more diffuse and hematopoietically diverse nature of bone marrow, with significant fat replacement in adult animals. Yet, each contained comparable estimated total cell counts (**Table 1**), even with our selective marrow gating, due to the much larger total volume of marrow versus spleen, suggesting the marrow contained at least equivalent engrafted B cells/plasma cells.

### Examining infused B cell distribution within the marrow compartment

As noted above, there was considerable heterogeneity of labeled cell distribution, especially within the bone marrow compartment. Due to scan lengths and constraints on anesthesia time, PET imaging of RMs is usually limited to a region spanning from the cervical spine/skull base down to the distal femurs/patellae. For the initial and repeat infusions into Subject 2, we were able to perform full body scans on day +1, when the bone marrow signal is most robust due to minimal radioisotope decay, allowing reasonable scan times. These full body scans allowed collection of data from the distal extremities, skull, and coccygeal (tail) spine (**Figure 5A**, **Supplemental Table 3**).

Both visual inspection and quantitation of ^89^Zr activity on the PET images demonstrated that the marrow signal is primarily confined to the proximal limbs and axial skeleton. Indeed, the narrower scans performed at the other timepoints only excluded approximately 10% of the total signal, indicating that they were sufficient to capture density and localization trends. Within the skeleton, approximately 60% of activity was in the pelvis and spine, primarily the thoracolumbar and sacral regions, with smaller contributions from the cervical spine, and effectively none from the tail. In the extremities, the activity was confined to the proximal long bones (humeri and femurs) with minimal contribution from the forearms and lower legs, and no detectible activity in the hands. The skull had a diffuse, inhomogeneous distribution with an apparent preference towards the skull base compared to the calvarium.

Even within areas with the most signal, such as the spine, the bone marrow examined more closely by overlaid PET/CT shows further inhomogeneities. The PET signal demonstrated a punctate distribution with intervening areas of diminished or absent activity (**Figure 5B**, **Supplemental Figure 4**), creating discontinuities in signal within the marrow space. Although some gaps could be explained by the segmental nature of vertebral bodies and intervening vertebral discs, even within the long bones there were areas of discontinuity. Also, the foci of increased intensity did not appear to correlate between the initial infusion and the re-infusion in Animal #2 (**Figure 5C**, **Supplemental Figure 4**). This heterogeneity suggests particular marrow neighborhoods conducive to supporting infused B cells/plasma cells. To further examine this inhomogeneity, we conducted a spine MRI on the Subject 2 immediately prior to the second cell infusion (**Figure 5C**). Neither STIR that suppresses intensity of fat nor T1-weighted sequences were able to clearly correlate the distribution of the PET signal with the distribution of red (active) or yellow (adipose) marrow, although MRI may be of insufficient resolution to resolve more localized structures.

## Discussion

The dynamics and localization of cellular therapy products following infusion are critical data for improving the function of such products. Stable homing to physiologically appropriate tissues is essential to the long-term engraftment of cell products; whereas homing to disadvantageous locations can lead to graft failure through either acute loss or reduced cellular lifespan – both being important considerations for therapies intended as alternatives to infusion-based biologics delivery. The techniques we present here allow for non-invasive, real-time monitoring of engraftment of a disseminated population of B cells at sub-organ resolution using high sensitivity PET imaging.

We have demonstrated that ex vivo manipulated and expanded RM B cells can be differentiated towards more mature plasma cells and reinfused into an autologous animal as proof-of-concept relevant to the development of B cell medicines. Following ex vivo expansion from blood B cells, ^89^Zr-oxine labeling, and infusion, the early signal seen within the lungs on PET/CT images was expected, as the entire venous return passes through the pulmonary vasculature prior to returning to systemic circulation. Consequently, the pulmonary capillary beds, which already house significant immune tissues, readily provide a place for the cells to transiently linger. Approximately 20-30% of the infused cells then localized to the spleen and bone marrow, physiologic niches for both B cell and plasma cells. As the main lymphoid organ of the body and the primary reservoir of extravascular lymphocytes, it is not surprising that the spleen has a higher density of infused cells compared to marrow. By the end of the 6-day observation period, the cells in these locations remained stable as a fraction of the total ^89^Zr activity remaining in the body, suggesting stable homing and lodgment. Monitoring cell engraftment long-term will require developing RM-specific engineering protocols using reporter systems such as the sodium iodide symporter (NIS) or non-imaging-based tracking technologies.^46^

**Figure 2.**
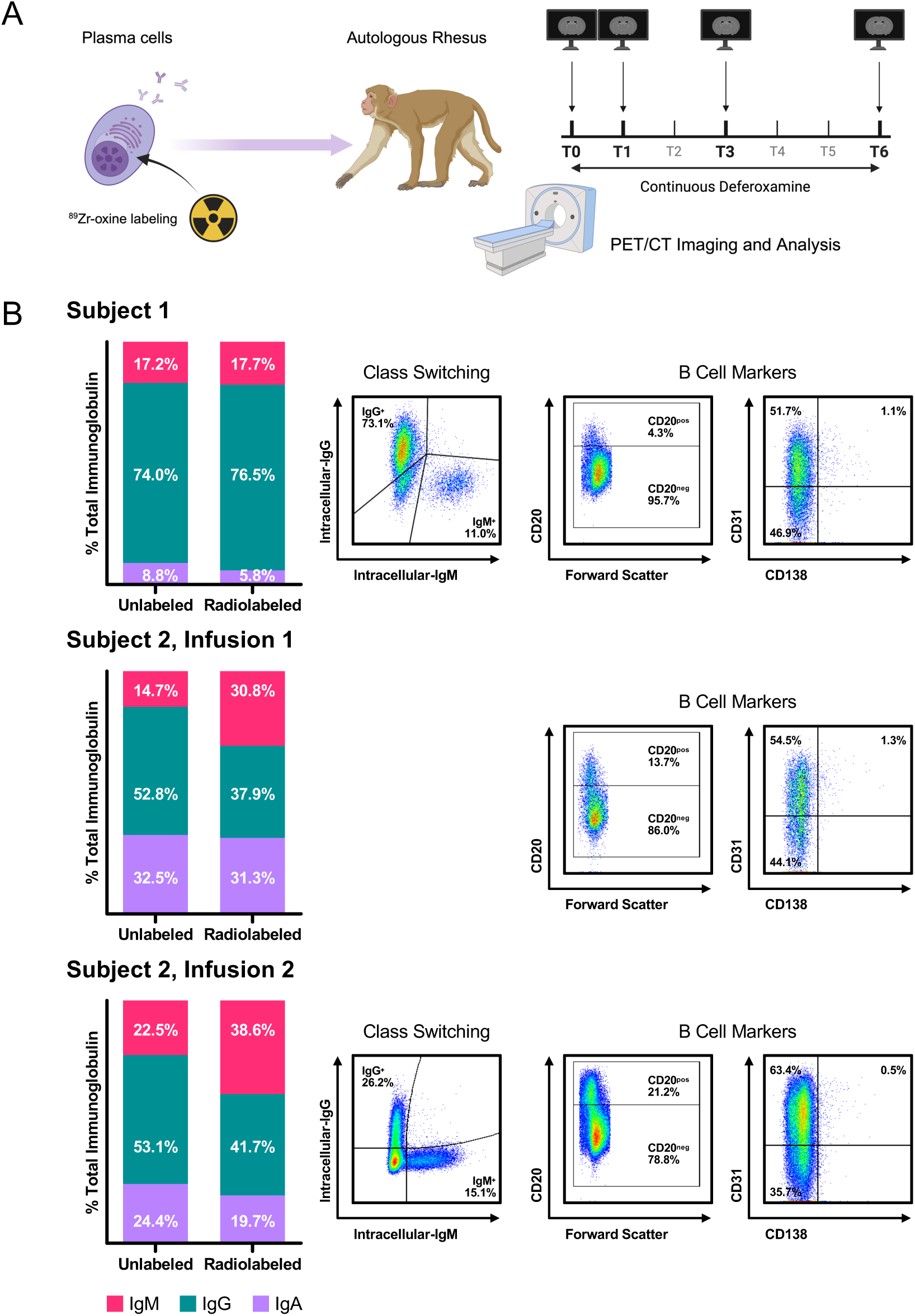
^89^Zr-oxine labeling of rhesus macaque B cells. A. Experimental schema for the expansion, ^89^Zr-oxine labeling, infusion, and post-infusion monitoring of autologous non-human primate expanded and differentiated B cells. B. Analyses were conducted on the day of infusion for each of the three cell products. ELISpot analysis (left) demonstrates the production of IgM, IgG and IgA in cells with and without ^89^Zr-oxine labeling. Flow cytometry (right) demonstrates intracellular immunoglobulin, CD20, CD31 and CD138 expression. Immunoglobulin flow cytometry for subject 2, infusion 1 not included due to technical failure.

**Figure 3.**
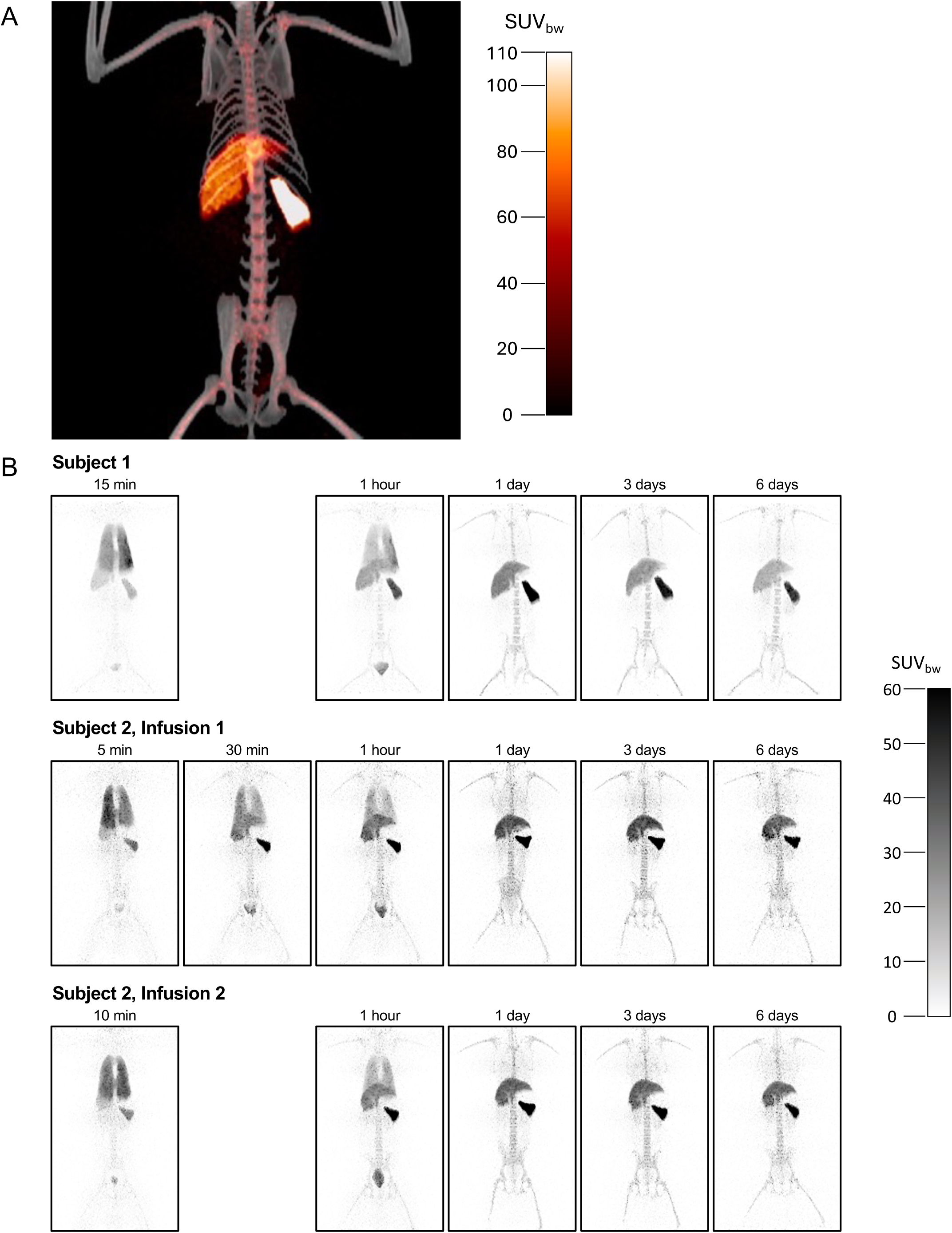
Infusion of autologous ^89^Zr-oxine-labeled rhesus macaque B cells. A. Representative day +1 PET image (orange) merged with CT image (grayscale) from Subject 1. B. Sequential PET imaging of Subject 1 and 2 after infusion of *ex vivo* expanded, ^89^Zr-oxine-labeled autologous B cells (top two rows) and on Subject 2 following a second infusion 70 days later (bottom row). Each frame shows the anteroposterior maximum intensity projection images shaded according to the standardized uptake value (SUV) at the indicated timepoints.

**Figure 4.**
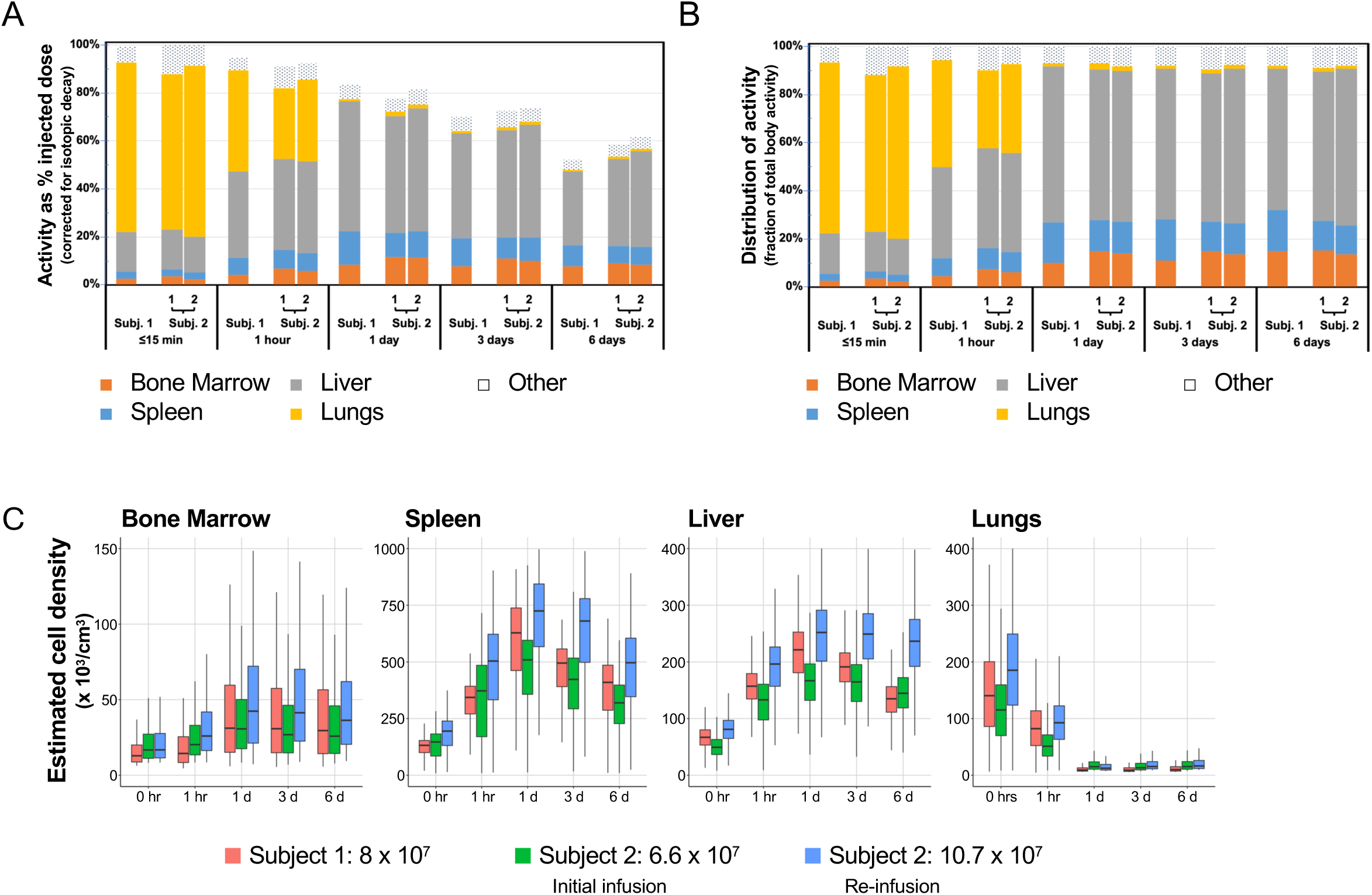
Quantitation of the distribution and persistence of *ex vivo* expanded rhesus macaque B cells in various organs following infusion. Volumes-of-interest around the indicated target organs were quantitated for their ^89^Zr activity after correcting for radioisotope decay. A. Total activity within the indicated organs as a percentage of the original infused dose is plotted according to time. B. The activity of each organ of interest as a fraction of the total body activity remaining is plotted as a function of time for each animal and each infusion. C. Specific activity (mBq/cell) corrected for decay was used to determine cell density per unit volume in key organs and the distributions of intensities plotted as whisker box plots according timepoint for each infusion.

**Figure 5.**
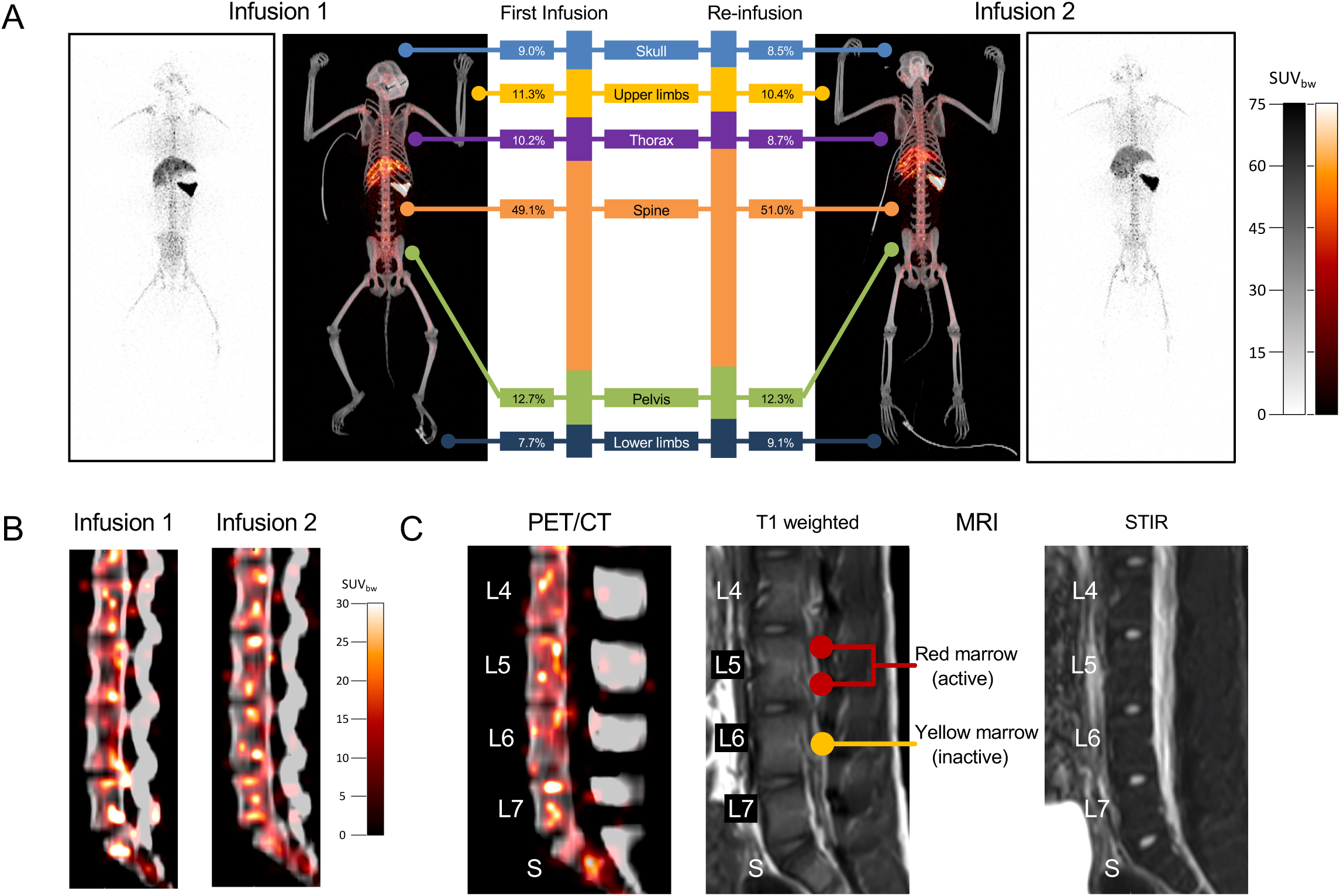
Marrow distribution of *ex vivo* expanded non-human primate B cells following re-infusion. A. Whole-body PET/CT imaging was conducted on day +1 for the initial infusion (left) and reinfusion (right) of ^89^Zr-oxine labeled B cells in Subject 2. The activity of the entire skeleton was divided according to anatomic regions, the percentage contribution of each is shown. B. Representative, corresponding sagittal slices of the lumbosacral spine demonstrating foci of increased activity (local signal maxima) from each infusion for Subject 2; initial infusion (left) and reinfusion (right). C. Prior to the reinfusion, MRI was performed on the spine of subject 2. Representative corresponding sagittal slices from the PET/CT and pre reinfusion MRI imaging for Subject 2 are shown. The approximate location of red (active) and yellow (adipose) marrow as indicated by STIR and T1 intensity are shown.

**Table 1.** Minimum cell occupancy of each organ as estimated from total organ, decay-adjusted ^89^Zr activity (percent of infused dose).

**Subject 1: $7.97 \times 10^7$ cells**
|  | Bone Marrow | Spleen | Liver | Lungs | Whole Body |
| --- | --- | --- | --- | --- | --- |
| 15 min | $2.1 \times 10^6$ (2.7%) | $2.2 \times 10^6$ (2.8%) | $1.3 \times 10^7$ (16.7%) | $5.6 \times 10^7$ (70.7%) | $7.9 \times 10^7$ (99.6%) |
| 1 hour | $3.3 \times 10^6$ (4.1%) | $5.7 \times 10^6$ (7.1%) | $2.9 \times 10^7$ (36.0%) | $3.4 \times 10^7$ (42.2%) | $7.5 \times 10^7$ (94.8%) |
| 1 day | $6.6 \times 10^6$ (8.2%) | $1.1 \times 10^7$ (14.2%) | $4.3 \times 10^7$ (54.0%) | $8.6 \times 10^5$ (1.2%) | $6.6 \times 10^7$ (83.3%) |
| 3 days | $6.2 \times 10^6$ (7.8%) | $9.4 \times 10^6$ (11.8%) | $3.5 \times 10^7$ (43.7%) | $7.8 \times 10^5$ (1.0%) | $5.6 \times 10^7$ (69.9%) |
| 6 days | $6.2 \times 10^6$ (7.8%) | $7.0 \times 10^6$ (8.8%) | $2.4 \times 10^7$ (30.6%) | $4.6 \times 10^5$ (0.6%) | $4.1 \times 10^7$ (52.0%) |

**Subject 2: $6.64 \times 10^7$ cells**
|  | Bone Marrow | Spleen | Liver | Lungs | Whole Body |
| --- | --- | --- | --- | --- | --- |
| 5 min | $2.3 \times 10^6$ (3.4%) | $2.0 \times 10^6$ (3.1%) | $1.1 \times 10^7$ (16.3%) | $4.3 \times 10^7$ (65.0%) | $6.6 \times 10^7$ (99.7%) |
| 30 min | $3.7 \times 10^6$ (5.6%) | $4.3 \times 10^6$ (6.5%) | $2.1 \times 10^7$ (31.9%) | $2.6 \times 10^7$ (39.7%) | $6.3 \times 10^7$ (94.5%) |
| 1 hour | $4.4 \times 10^6$ (6.7%) | $5.3 \times 10^6$ (8.0%) | $2.5 \times 10^7$ (37.9%) | $1.9 \times 10^7$ (29.2%) | $6.0 \times 10^7$ (91.0%) |
| 1 day | $7.7 \times 10^6$ (11.6%) | $6.7 \times 10^6$ (10.0%) | $3.2 \times 10^7$ (48.8%) | $1.3 \times 10^6$ (1.9%) | $5.2 \times 10^7$ (77.8%) |
| 3 days | $7.2 \times 10^6$ (10.8%) | $6.0 \times 10^6$ (9.0%) | $3.0 \times 10^7$ (44.6%) | $8.1 \times 10^5$ (1.2%) | $4.8 \times 10^7$ (72.7%) |
| 6 days | $5.9 \times 10^6$ (8.9%) | $4.8 \times 10^6$ (7.2%) | $2.4 \times 10^7$ (36.3%) | $6.1 \times 10^5$ (0.9%) | $3.9 \times 10^7$ (58.5%) |

**Subject 2 (re-infusion): $1.07 \times 10^8$ cells**
|  | Bone Marrow | Spleen | Liver | Lungs | Whole Body |
| --- | --- | --- | --- | --- | --- |
| 10 min | $2.3 \times 10^6$ (2.2%) | $3.1 \times 10^6$ (2.9%) | $1.6 \times 10^7$ (14.9%) | $7.6 \times 10^7$ (71.4%) | $1.1 \times 10^8$ (99.7%) |
| 1 hour | $6.2 \times 10^6$ (5.8%) | $8.2 \times 10^6$ (7.6%) | $4.1 \times 10^7$ (38.0%) | $3.6 \times 10^7$ (34.0%) | $9.8 \times 10^7$ (92.2%) |
| 1 day | $1.2 \times 10^7$ (11.3%) | $1.2 \times 10^7$ (11.0%) | $5.5 \times 10^7$ (51.1%) | $1.7 \times 10^6$ (1.6%) | $8.7 \times 10^7$ (81.7%) |
| 3 days | $1.1 \times 10^7$ (10.0%) | $1.0 \times 10^7$ (9.6%) | $5.0 \times 10^7$ (47.1%) | $1.2 \times 10^6$ (1.1%) | $7.8 \times 10^7$ (73.5%) |
| 6 days | $9.0 \times 10^6$ (8.4%) | $7.9 \times 10^6$ (7.4%) | $4.3 \times 10^7$ (40.1%) | $9.0 \times 10^5$ (0.8%) | $6.6 \times 10^7$ (61.6%) |

**Table 2.** Total organ occupancy as a percentage of total detected cells at each time point.

**Subject 1**
|  | **Bone Marrow** | **Spleen** | **Liver** | **Lungs** |
| --- | --- | --- | --- | --- |
| 15 min | 2.7% | 2.8% | 16.7% | 71.0% |
| 1 hour | 4.4% | 7.5% | 38.0% | 44.6% |
| 1 day | 9.9% | 17.1% | 64.8% | 1.3% |
| 3 days | 11.1% | 16.9% | 62.6% | 1.4% |
| 6 days | 15.0% | 17.0% | 58.8% | 1.1% |

**Subject 2**
|  | **Bone Marrow** | **Spleen** | **Liver** | **Lungs** |
| --- | --- | --- | --- | --- |
| 5 min | 3.4% | 3.1% | 16.4% | 65.2% |
| 30 min | 5.9% | 6.9% | 33.7% | 42.0% |
| 1 hour | 7.3% | 8.8% | 41.6% | 32.1% |
| 1 day | 14.9% | 12.9% | 62.7% | 2.4% |
| 3 days | 14.9% | 12.4% | 61.4% | 1.7% |
| 6 days | 15.3% | 12.3% | 62.0% | 1.6% |

**Subject 2 (re-infusion)**
|  | **Bone Marrow** | **Spleen** | **Liver** | **Lungs** |
| --- | --- | --- | --- | --- |
| 10 min | 2.2% | 2.9% | 14.9% | 71.7% |
| 1 hour | 6.3% | 8.3% | 41.2% | 36.8% |
| 1 day | 13.8% | 13.5% | 62.6% | 2.0% |
| 3 days | 13.6% | 13.1% | 64.1% | 1.5% |
| 6 days | 13.7% | 12.0% | 65.1% | 1.4% |

**Table 3.** Estimated median organ cell density x 10^3^ cells/cm^3^ (range).

**Subject 1**
|  | **Bone Marrow** | **Spleen** | **Liver** | **Lungs** |
| --- | --- | --- | --- | --- |
| 15 min | 12.89 (6.3-105.8) | 132.01 (13.4-248.8) | 66.95 (6.6-168.7) | 141.77 (6.4-661.2) |
| 1 hour | 14.42 (4.6-189.5) | 343.41 (45.4-537.4) | 157.15 (9.5-293.3) | 82.22 (4.6-397.9) |
| 1 day | 31.67 (5.9-315.2) | 628.27 (109.1-909.4) | 221.39 (5.9-363.6) | 9.21 (5.9-73.6) |
| 3 days | 30.12 (5.5-280.9) | 495.01 (66.3-687.1) | 191.26 (11.7-333.3) | 8.87 (5.5-70.1) |
| 6 days | 37.41 (5.7-292.4) | 409.82 (10.3-691.5) | 134.92 (7.8-302.9) | 9.93 (5.7-79.2) |

**Subject 2, Infusion 1**
|  | **Bone Marrow** | **Spleen** | **Liver** | **Lungs** |
| --- | --- | --- | --- | --- |
| 5 min | 16.76 (8.2-171.2) | 146.90 (8.3-344.2) | 49.34 (8.2-166.4) | 115.29 (8.3-452.7) |
| 30 min | 18.53 (8.3-226.0) | 315.00 (8.9-579.5) | 108.13 (8.5-420.1) | 71.28 (8.4-246.9) |
| 1 hour | 20.27 (8.5-285.7) | 372.29 (9.3-716.0) | 133.09 (8.8-344.1) | 51.14 (8.5-294.2) |
| 1 day | 31.03 (8.3-320.5) | 509.13 (12.5-926.5) | 166.74 (8.4-334.7) | 15.01 (8.3-133.7) |
| 3 days | 27.11 (7.0-386.4) | 422.80 (16.5-809.1) | 164.76 (7.3-371.9) | 13.10 (7.0-118.4) |
| 6 days | 26.23 (7.8-297.1) | 319.35 (9.7-595.7) | 144.82 (9.1-563.7) | 15.28 (7.8-95.5) |

**Subject 2, Infusion 2**
|  | **Bone Marrow** | **Spleen** | **Liver** | **Lungs** |
| --- | --- | --- | --- | --- |
| 10 min | 16.81 (8.3-150.5) | 194.76 (13.8-435.5) | 81.03 (8.4-255.1) | 185.83 (8.4-458.5) |
| 1 hour | 26.06 (8.5-305.0) | 503.99 (11.8-903.7) | 196.29 (8.5-389.7) | 92.62 (8.5-342.8) |
| 1 day | 44.07 (7.3-389.0) | 733.29 (176.8-1151.2) | 252.36 (8.2-486.4) | 12.38 (7.3-192.1) |
| 3 days | 42.80 (8.8-405.0) | 684.30 (15.0-1116.5) | 249.38 (9.1-569.2) | 15.35 (8.8-101.5) |
| 6 days | 37.37 (9.4-479.1) | 496.56 (24.0-891.4) | 236.79 (10.2-506.0) | 16.39 (9.4-106.3) |

Several interesting observations regarding the pattern of localization of cells in the bone marrow are of note. First, although signal can be identified in almost every part of the skeleton containing marrow, the pattern is not even throughout. Despite encompassing almost half of the skeleton, less than a fifth of the signal is localized to the bones in the extremities. Similarly, the skull, one-fifth of the skeleton, has less than a tenth of the signal. In contrast, the spine and pelvis, less than a third of the skeleton, contained almost two-thirds of the entire ^89^Zr activity, 95% of which was further confined to the thoracolumbar spine and pelvis. Indeed, even within the extremities, over 90% of the activity was confined to the proximal long bones, and with further preference for the proximal ends of these bones. Although this imaging monitors infused B cells, prior work indicates that there is considerable overlap between the HSC and B lymphocyte niches.^47^ Thus, B cell imaging may provide insights into the hematopoietic niche, in general.

The axial and proximal distribution of the bone marrow signal, as opposed to a more global, uniform distribution, mirrors the distribution of active hematopoiesis within the human bone marrow space, especially within adolescents and adults.^48–50^ This contrasts with the distribution of active bone marrow in mice which is more uniformly distributed throughout the skeleton.^51–54^ In mice, intravital imaging has demonstrated that within the femur, hematopoietic cells distribute almost uniformly along the entire diaphysis.^53,54^ Furthermore, mouse tibiae are rich sources of hematopoietic cells.^51,52^ The pattern we observed suggests that NHP bone marrow architecture and hematopoietic cellularity better reflects the distribution of active human hematopoiesis than the mouse, further underscoring the difference between murine and primate hematopoiesis and bone marrow architecture.

Beyond the contraction of active niches supporting infused B lineage cells to the axial skeleton, there is further heterogeneity in the ^89^Zr activity distributed in the marrow, most evident in the vertebral bodies. Some of the marrow signal heterogeneity likely reflects the stochastic nature of detecting the relatively low signals therein; however, the marrow signal also contains foci crossing multiple imaging planes with intensities comparable to that of the spleen, consistent with actual clusters of infused labeled cells. On bone marrow biopsies, lymphoid cells can present as either diffusely scattered throughout the active marrow space or in lymphoid aggregates including B cell aggregates.^55–57^ Our imaging findings suggest that the infused cells may be sub-localizing to specific areas of the marrow rich in supportive niches. Finally, in the animal receiving a second infusion, the localization of foci did not match between infusions, and the degree of marrow homing was similar, suggesting that doses of up to ten million cells per kilogram did not “fill” marrow or spleen niches, leaving remaining “space”, relevant for clinical applications.

An unexpected finding was the large and sustained hepatic signal shown on the PET images. The liver receives approximately 25% of all cardiac output, and thus it is not surprising that within a few hours of circulation, especially as the cells begin to mobilize from the pulmonary system, significant portions will transit through the liver. Yet, in comparison with the pulmonary signal, the hepatic signal was persistent: the decay-corrected signal intensity only decreasing with a 1-2-week half-life. There are several possible interpretations. The liver signal may be artifactual, representing cell-free (but possibly protein-bound) ^89^Zr that becomes sequestered in the liver from dying cells. However, continuous infusion of deferoxamine ensures that label released by cell death or infused unbound ^89^Zr-oxine, if any, will be incorporated into ^89^Zr-deferoxamine complexes. Although deferoxamine complexes may be metabolized in the liver, elimination is primarily renal, as reflected by the signal found within the bladder. Indeed, the relatively low signal within the renal collection system and bladder indicate low levels of free ^89^Zr following infusion. Furthermore, deferoxamine complexes, most notably ferrioxamine, have terminal clearance half-lives of approximately 3-6 hours.^58,59^ That the liver signal declines with kinetics 20- to 50-fold slower indicates that this signal represents labeled cells. The lack of non-specific signal elsewhere in large tissues and organs (i.e., musculature, cortical bone, brain, etc.) and incapability of free ^89^Zr to label cells, argues against a non-specific labeling of tissues, leaving infused cells as the probable source of the signal. Although cells will encounter liver sinusoids after leaving the pulmonary circulation and then the heart, for up to two-thirds of the signal to end in the liver suggests a non-random engraftment process, either through selective homing of the cells to the liver or entrapment of cells by the liver as they transit the sinusoids.

The presence of immune cells within the liver is well-established, for example, most of the macrophages in the body can be found within the liver, as well as a large fraction of natural killer cells and other innate lymphoid cells, with B cells comprising up to 8% of non-hepatocytes.^60–62^ The liver microenvironment contains a medley of cytokines, even when healthy, including several important to B cell development, notably interleukin-7 (IL-7).^63–66^ Indeed, B cell recruitment to the liver is an important component of the immune response to bacterial and parasitic infections. Hepatic B cell localization has also been seen in disease processes such as precursor B cell lymphoblastic leukemia (over 40% of pediatric and adult cases present with hepatomegaly).^67^ All of these findings suggest that the liver may support B cell survival and expansion. Importantly, although B cells have been implicated in certain liver pathologies, B cell infiltration can be asymptomatic (e.g., B-ALL) or even beneficial (e.g., clearance and recovery from *Ehrlichia*),^68^ indicating that B cell localization itself is likely not damaging to the liver, an important consideration for B cell medicine development. The macaques receiving ^89^Zr-oxine labeled cells in our study showed only very mild increases in liver transaminases, with values remaining within normal limits on all but one determination and decreasing to baseline while signal persisted in the liver. There was no evidence of liver dysfunction, with stable albumin. The pattern of changes seen was instead consistent with the prolonged anesthesia required to perform imaging.

We conclude that these short-term tracking studies suggest that viable ex vivo generated B cell medicines (e.g., plasma cells and plasmablasts) are able to home to native environments such as the spleen and marrow, and stably persist for at least 6 days, without evident toxicity, encouraging for clinical development. Remarkably, they are able to engraft without any conditioning of the animals and can be equally efficiently readministered as needed. Important questions remain. Do cells that home to the liver persist there long term, redistribute to other tissues, or die out due to insufficient microenvironmental support? Can these hepatic B cells contribute to effective B cell medicine therapeutic effects? Do different B cell lineage cells or plasma cell subpopulations preferentially home to the spleen and marrow versus the liver? Answering these questions will advance both our understanding of B cell biology and the emerging field of B cell engineering.

## Materials and Methods

### Animals

All animals were housed and handled in accordance with protocols approved by the Animal Care and Use Committee (ACUC) of the National Heart, Lung, and Blood Institute. Animals were housed in specific pathogen free colonies and had not been previously used for experimentation. All procedures were conducted in accordance with good veterinary practice. All reagents used for in vivo purposes were of pharmaceutical grade except ^89^Zr-oxine, which was prepared sterilely.

### Cell Collection and Expansion

All cells were collected without mobilization. For small-scale *in vitro* testing, whole blood was collected via venipuncture. For large-scale experiments and infusion product generation, cells were collected via apheresis performed with either a CS3000 Plus (Baxter-Fenwal) or an Amicus Separator (Fresenius Kabi), primed using allogeneic, leukoreduced RM blood.^69^ Mononuclear cells were isolated using Ficoll-Paque PLUS (Cytiva). B cells were labeled with anti-non-human primate CD20 MicroBeads (Miltenyi) according to manufacturer protocols, and isolated by positive magnetic selection (LS columns and QuadroMACS Separator, Miltenyi). The isolated B cells were characterized using flow cytometry for CD20-positivity to ensure >90% purity. Cells were either expanded fresh or stored viably in CryoStor CS10 (StemCell Technologies) in vapor phase liquid nitrogen. All cells infused into animals were first frozen viably after selection to allow time for animal preparation.

Isolated B cells were plated in ImmunoCult-XF B Cell Base medium supplemented with Human B Cell Expansion Supplement (StemCell Technology), seeding at 2.5 x 10^5^ cells/mL in an appropriate format to provide media at a depth of 2-5 mm. Additional media was added without replacement every 2-3 days to maintain a density of 2.5 x 10^5^ cells/mL, using gentle trituration to mix and disrupt any aggregates. Cells were collected after 7 days of expansion for injection, experimentation, or cryopreservation.

For in vitro plasma cell differentiation, day 7 expanded cells were counted, washed and replated at a concentration of 2.5 x 10^5^ cells/mL in ImmunoCult-XF supplemented with 50 ng/mL human IL-2, 50 ng/mL IL-6, 50 ng/mL IL-10, and 10 ng/mL IL-15 and cultured for an additional 3 days. On Day 10 differentiated cells were counted, washed, and replated at a concentration of 2.5 x 10^5^ cells/mL in ImmunoCult-XF supplemented 50 ng/mL IL-6, 10 ng/mL IL-15, and 15 ng/mL IFN-α 2b (PBL Assay Sci, all other cytokines from Peprotech) and cultured for an additional 2 days.

### Flow Cytometric Analysis

Cell purity and B cell immunophenotyping were determined by multicolor flow cytometric analysis on either an LSRFortessa (BD Biosciences) or NovoCyte-Penteon (Agilent) cell analyzer. Live-dead discrimination was performed using the Live/Dead Fixable Near-IR Dead Cell Stain Kit (Invitrogen). Cells were blocked with Human TruStain FcX (BioLegend) and stained using the listed antibodies (**Supplemental Table 1**). For immunophenotyping, the cells were further treated with IC fixation and permeabilization buffers (Invitrogen), and stained with anti-IgG and anti-IgM antibodies (**Supplemental Table 1**). Purity and plasma cell staining master-mixes were created in Flow Cytometry Staining Buffer (eBioscience), while intracellular staining master-mix was created in permeabilization buffer. Brilliant Stain Buffer (BD Biosciences) was added to all staining master-mixes. The data were analyzed using FlowJo (v10.10.0).

### ELISpot Analysis

At least one day prior to plating cells, sterile MultiScreen 96-well PVDF membrane-lined ELISpot plates (MilliporeSigma) were activated with 35% ethanol (Fisher Scientific), washed 3 times with DPBS (Gibco) and coated with 10 mcg/mL of primary antibodies (**Supplemental Table 2**) overnight at 4 °C. On the day of plating, plates were blocked with 10% fetal bovine serum (Gibco) in sterile IMDM (Gibco) for 2 hours. Cells to be analyzed were added in 5-fold dilutions ranging from 10,000 to 0.64 cells/well in duplicate in IMDM with 10% fetal bovine serum along with negative control (no cells). Cells were incubated overnight at 37°C, 5% CO_2_. Unbound cells were removed, the membranes were washed with 1X Tween20 (Thermo Scientific) in deionized water, blocked with 1% bovine serum albumin (Fisher Scientific), and the appropriate secondary biotinylated antibodies (**Supplemental Table 2**) added at a 1:1000 dilution. Antibody secreting cells were measured after washing with 1X Tween20, incubation with alkaline phosphatase-conjugated streptavidin (Invitrogen) in 1% bovine serum albumin, and detection with BCIP/NBT substate (Sigma-Aldrich) according to manufacturer instructions. Spots were visualized and quantified using an ELISpot plate reader (Autoimmun Diagnostika). Frequency of antibody secreting cells was calculated and analyzed using GraphPad Prism (Dotmatics).

### ^89^Zr-oxine Cell Labeling

^89^ZrCl_4_ was produced at the institutional cyclotron facility.^70^ It was reacted with oxine and neutralized, as previously described, to generate ^89^Zr-oxine complex.^40–42^ B cells were collected from culture and washed in serum-free PBS. All collections were performed by centrifugation for 10 minutes at 400 x *g* at room temperature. Cells were incubated with approximately 55.5 kBq ^89^Zr-oxine per million cells in serum-free PBS for 15 minutes at room temperature. The reaction was quenched using approximately ten volumes of 1% autologous RM serum in PBS. The cells were washed three times with 1% autologous serum in PBS to remove unbound ^89^Zr-oxine. The labeled cells were counted, and the specific activity was determined by a dose calibrator, with a target specific activity of 26-30 mBq/cell (approximately 50% labeling efficiency). The product was resuspended in approximately 7 mL pharmaceutical-grade isotonic saline with 1% autologous serum for injection (maximum density of 10^8^ cells/mL) or an appropriate volume of IMDM with 10% fetal bovine serum for downstream *in vitro* applications. Small aliquots of cells were reserved for flow cytometric (unlabeled) and ELISpot (both labeled and unlabeled) analyses.

### Cell Infusion

Beginning 10 minutes prior to cell infusion, 1-gram of deferoxamine was administered via central catheter at a rate of 15 mg/kg/hour. The animal was anesthetized, and the labeled cells were delivered by intravenous push followed by a saline flush. The activity of the syringe, including needle and cap, was measured by a bedside dose calibrator immediately prior to and following injection to determine final cell dose administered and the total activity administered for decay correction. Following the bolus of deferoxamine (approximately 5-10 hours), a maintenance infusion was initiated at a rate of 40 mg/kg/day, continuing through day +6 imaging. If intravenous access was lost during maintenance infusion, 500 mg deferoxamine was administered intramuscularly, repeating every 12 hours until completion of the experiment (Subject 1, after day +1).

### Image Analysis and Statistical Analysis

PET/CT images were performed on a Discovery MI DR (GE Healthcare) imaging system, acquiring for 5 minutes per bed position on Day 0, extending to 6, 8, and 12 minute on Day 1, 3 and 6, respectively, and 4 or 5 bed positions to cover the torso of the subject. In addition, Subject 2 underwent whole-body PET imaging on Day 1 at 2 minutes per bed, 8 bed positions. PET images were reconstructed using GE’s Q CLEAR algorithm. Analyses and visualization of acquired PET images were conducted with MIM software (v7.2.9). Volumes-of-interest (VOI) were drawn according to anatomic boundaries defined by CT images, subtracting any PET signal spillover from adjacent organs. To capture the total cell occupancy within a tissue, VOI contour was drawn to incorporate the entire PET signal associated with the tissue of interest. Activity was decay-corrected using the ^89^Zr half-life of 78.41 hours (282,276 sec). For cell density determination within organs (especially the marrow), a more parsimonious VOI was used that only incorporated the union of the CT-defined anatomical region and PET signal (see **Supplemental Figure 5** for examples). MRI were acquired on a 3T Ingenia Elition X (Philips) system using T1-weighted and short-T1 inversion recovery (STIR) sequences. All images were either analyzed with MIM software or were passed through to R for further statistical analyses.^71^ Within R, DICOM data processing was conducted with the RadOnc^72^ and oro.dicom^73^ packages. Visualization was conducted with the ggplot2 R package.^74^

## Data Availability Statement

Data presented in this paper are available upon request to David Young.

## Acknowledgments

The authors wish to acknowledge the technical support and excellent animal care provided by Allen Krouse, Seth Linde, Theresa Engels, Justin Golomb, NHLBI Animal Program Staff, and the NIH Division of Veterinary Resources. NHP apheresis processing was performed by Aylin Bonifacino. PET imaging was conducted by Mona Lisa Cedo, Joseph Carreon and Cristopher Leyson, and MRI by Dagane Daar and Hoa Tien, all of the Molecular Imaging Branch, NCI. Technical and logistical assistance was provided by Keyvan Keyvanfar, Stephanie Sellers, and Judy Yu of the NHLBI and Anja Hohmann, Lea Hachigian, and Cleo Isaacs of Be Biopharma. The graphical abstract and diagrams in **Figures 1A** and **2A** were created with BioRender.com. This work was supported through the NHLBI Division of Intramural Research, the NCI Division of Intramural Research, and in part through a Collaborative Research and Development Agreement (CRADA) with Be Biopharma (Cambridge, MA).

## Author Contributions

DJY, SH, NS, SA, ASL, RAM, and CED conceived of and designed studies. DJY, AJE, JT, EL, JB, SH, and KGQC conducted experiments. DJY, AJE, EL, KGQC, ASL, and NS conducted data analyses. DJY wrote the manuscript, and AJE and ASL contributed. All authors reviewed, edited, and approved the manuscript.

---

**Supplemental Figure 1.**
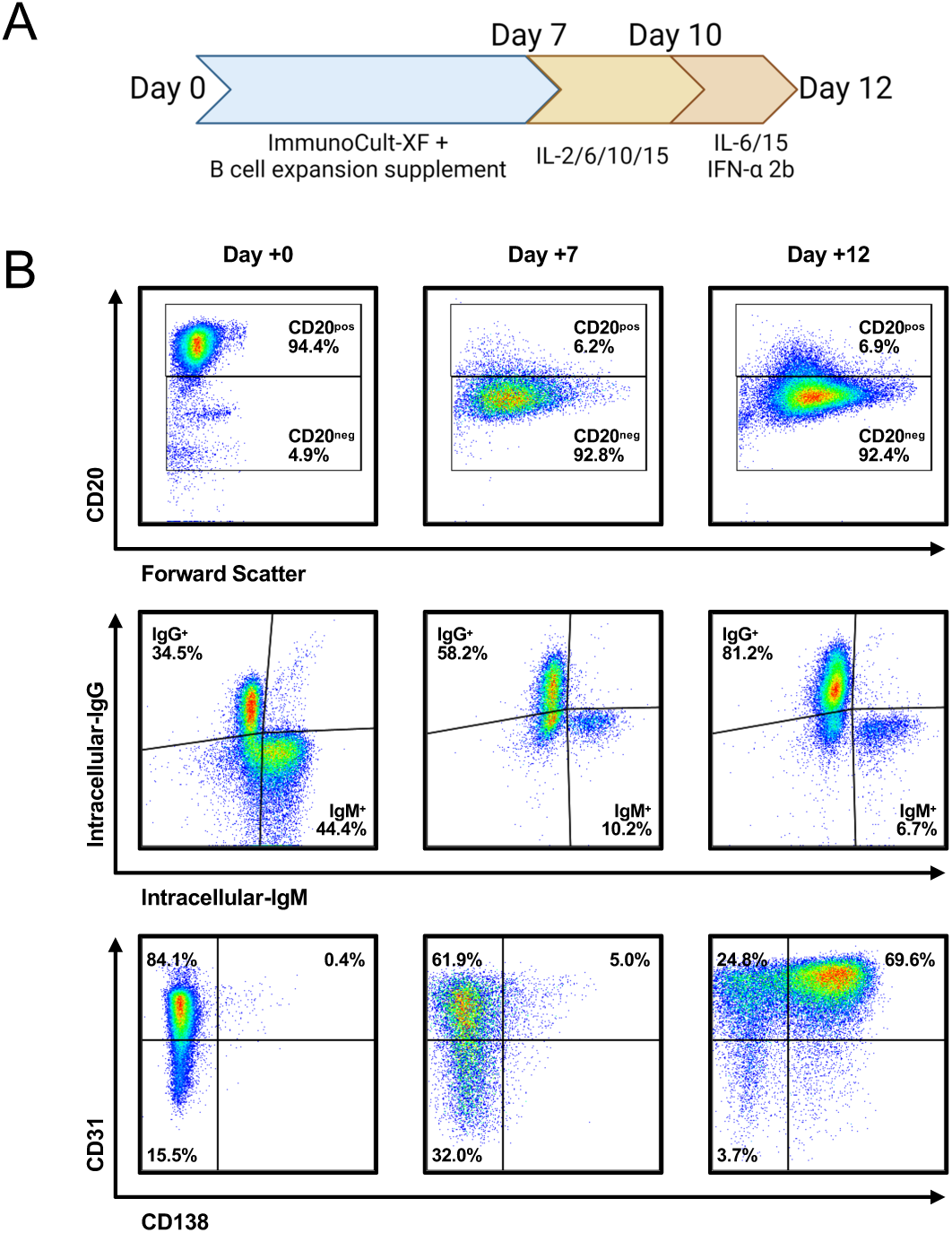
Analysis of ex vivo expanded B cells over culture time. A. Culture scheme for 12-day culture-based expansion. B. Extended flow cytometric analysis of NHP B cells during ex vivo expansion over approximately 12 days is shown with key B cell markers.

**Supplemental Figure 2.**
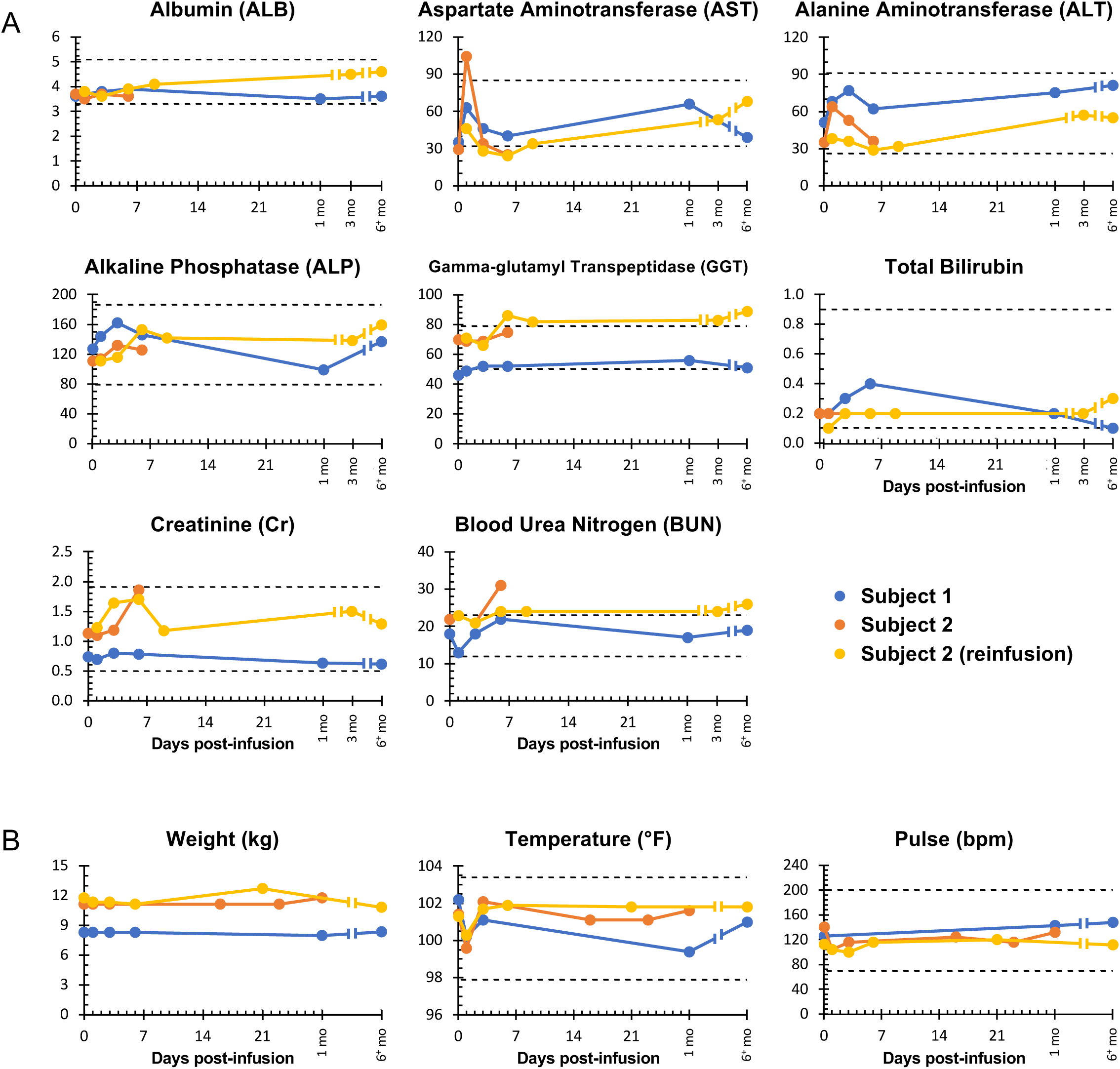
Clinical parameters during and following infusion of ^89^Zr-oxine-labeled rhesus macaque B cells. A. Laboratory values and B. clinical measurements were monitored during and following autologous infusion of *ex vivo* expanded ^89^Zr-oxine labeled non-human primate B cells. The indicated parameters are plotted by time from infusion (day 0). The re-infusion of subject 2 occurred 70 days following the initial infusion. Dashed lines indicate the upper and lower limits of the normal ranges for the two subjects.

**Supplemental Figure 3.**
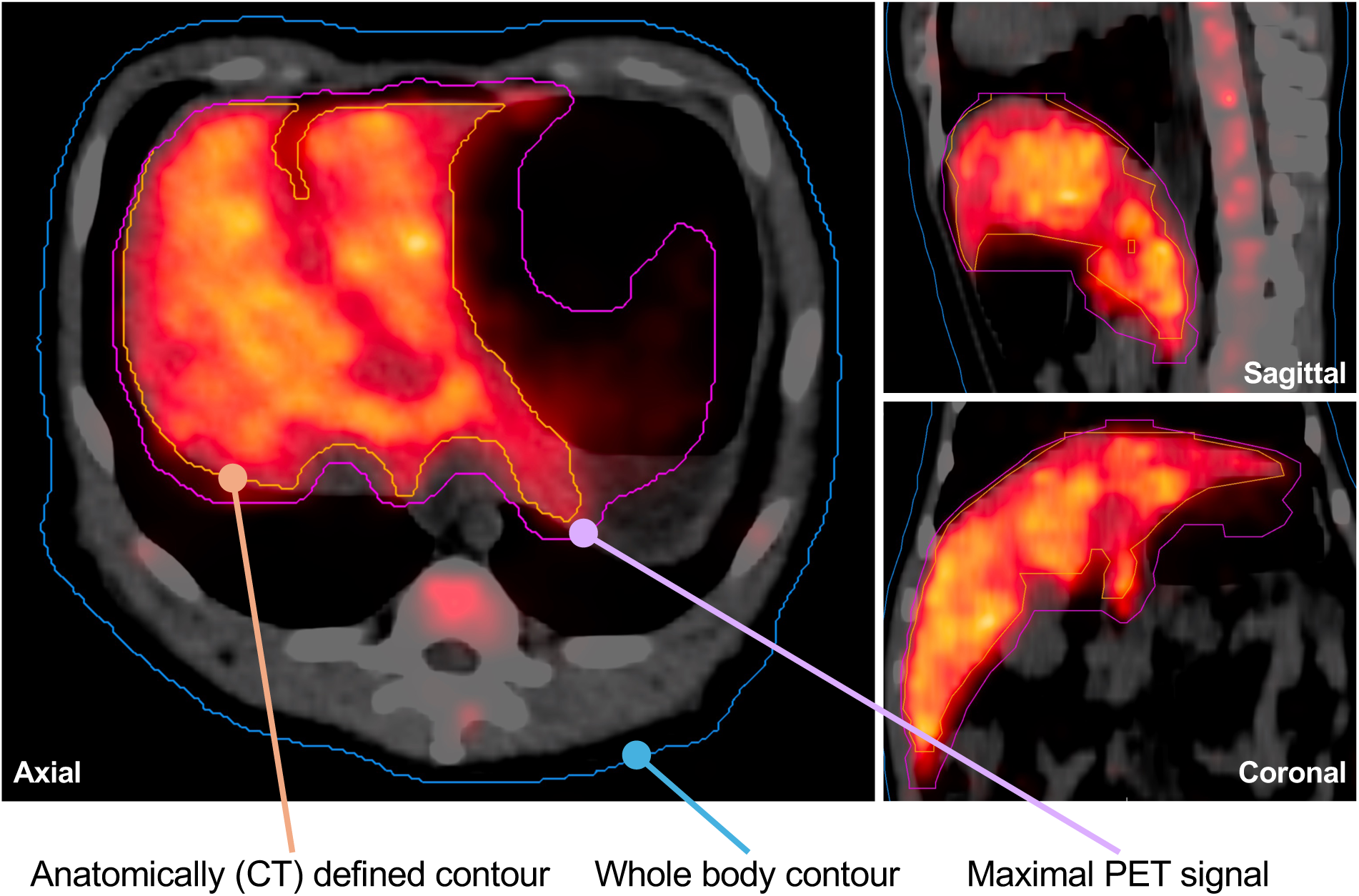
Construction of volumes-of-interest for image analysis. Representative axial (left), sagittal (top-right), and coronal (bottom-right) sections of PET/CT merged images showing the volumes-of-interest (VOI) contours of the liver. To calculate the total organ activity, a contour (magenta) was drawn on each PET/CT merged axial image slice to capture the entire anatomically defined region (by CT) as well as any additional PET signal that could be definitively attributed to the organ of interest (instead of adjacent organs). To calculate the cell density while minimizing irrelevant “dead space” (e.g., non-marrow bone, anatomic potential spaces, motion artifacts), a parsimonious contour (orange) was drawn to capture the union of the anatomic region that was also active on PET imaging. The whole-body contour (blue) represents the entire CT-defined body captured by the scan.

**Supplemental Figure 4.**
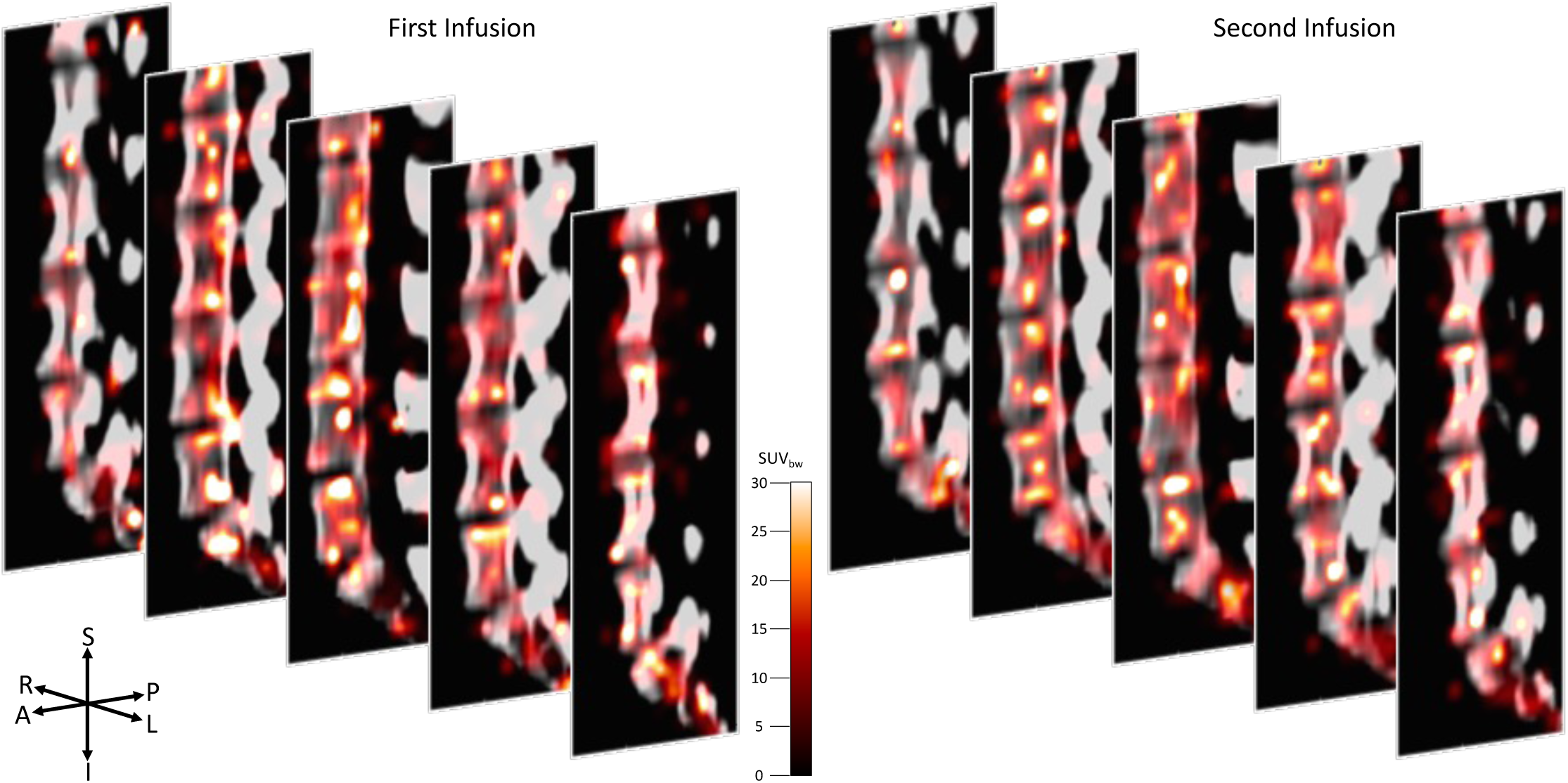
Comparison of the distribution of infused rhesus macaque B cells across different infusions. A series of paired sagittal slices from the lumbosacral spine comparing the distribution of ^89^Zr-oxine labeled B cells engrafted in the bone marrow foci on day +1 of infusion 1 and infusion 2 for Subject 2.

**Supplemental Table 1.** Antibodies used for flow cytometric analyses.

| Antigen | Conjugate | Manufacturer | Clone | Catalog | Use |
| --- | --- | --- | --- | --- | --- |
| CD3 | BV605 | BD Biosciences | SP34-2 | 562994 | 1,2 |
| CD14 | APC | Invitrogen | MEM-18 | MA1-10095 | 1 |
| CD20 | FITC | Biolegend | 2H7 | 302304 | 1,2 |
| CD56 | PE | Invitrogen | TULY56 | 12-0566-42 | 1 |
| CD184 | APC | Invitrogen | 12G5 | 17-9999-41 | 2 |
| CD14 | BV605 | Biolegend | M5E2 | 301834 | 2 |
| CD138 | PE | Invitrogen | DL-101 | 12-1389-42 | 2 |
| CD38 | mFluor450 | Caprico Biotech | OKT10 | 1008145 | 2 |
| CD31 | PE/Cy7 | Biolegend | WM59 | 303117 | 2 |
| IgG | BB700 | BD Biosciences | G18-145 | 742235 | 3 |
| IgM | BUV395 | BD Biosciences | G20-127 | 563903 | 3 |

1 Cell purity determination
2 Cell phenotype
3 Cell phenotype (intracellular staining)

**Supplemental Table 2.**
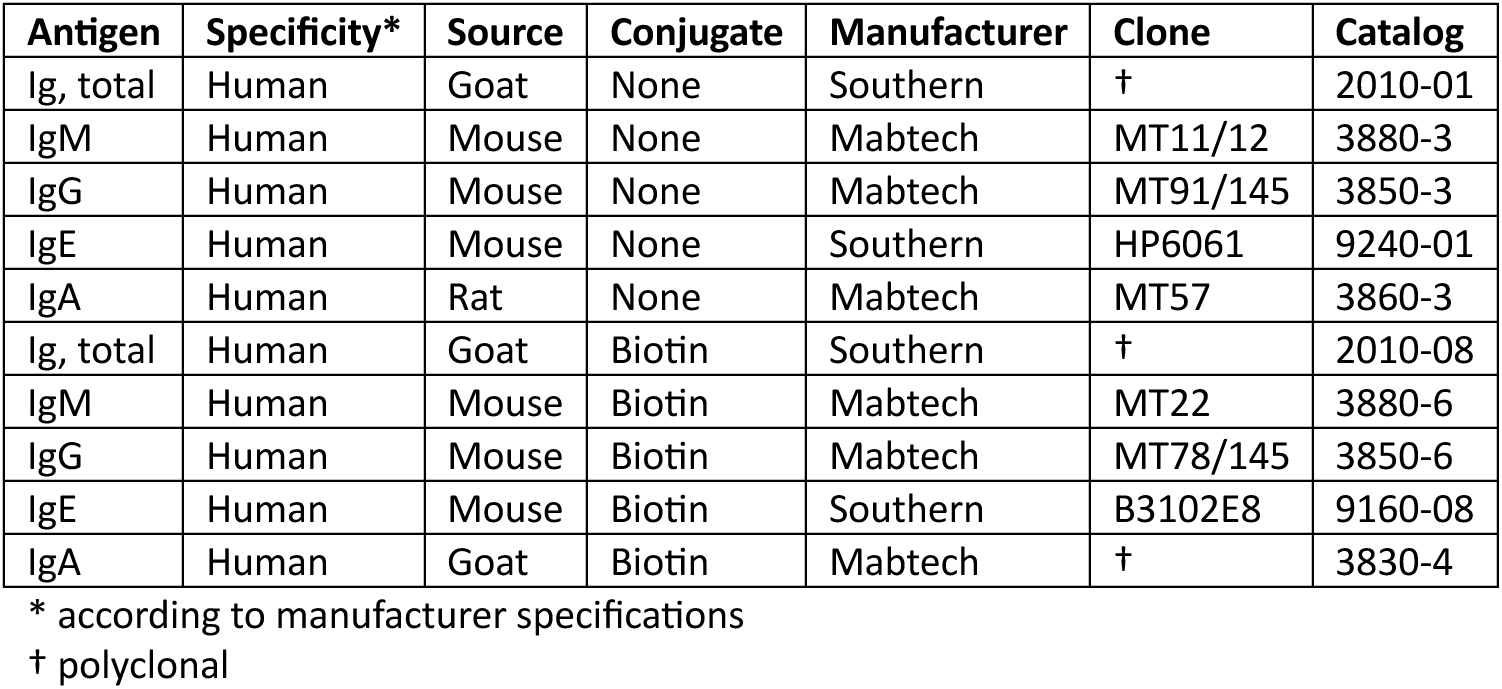
Antibodies used for ELISpot analyses.

| Antigen | Specificity* | Source | Conjugate | Manufacturer | Clone | Catalog |
| --- | --- | --- | --- | --- | --- | --- |
| Ig, total | Human | Goat | None | Southern | † | 2010-01 |
| IgM | Human | Mouse | None | Mabtech | MT11/12 | 3880-3 |
| IgG | Human | Mouse | None | Mabtech | MT91/145 | 3850-3 |
| IgE | Human | Mouse | None | Southern | HP6061 | 9240-01 |
| IgA | Human | Rat | None | Mabtech | MT57 | 3860-3 |
| Ig, total | Human | Goat | Biotin | Southern | † | 2010-08 |
| IgM | Human | Mouse | Biotin | Mabtech | MT22 | 3880-6 |
| IgG | Human | Mouse | Biotin | Mabtech | MT78/145 | 3850-6 |
| IgE | Human | Mouse | Biotin | Southern | B3102E8 | 9160-08 |
| IgA | Human | Goat | Biotin | Mabtech | † | 3830-4 |

\* according to manufacturer specifications
† polyclonal

**Supplemental Table 3.** Distribution of ^89^Zr activity throughout the skeleton.

| Region | Infusion 1 | Infusion 2 | Average | Volume<br>(% total skeleton) |
| --- | --- | --- | --- | --- |
| **Skull** | **9.04%** | **8.47%** | **8.77%** | **18.85%** |
| **Spine** | **49.09%** | **50.98%** | **49.99%** | **21.41%** |
| Cervical Spine | 3.04% | 3.18% | 3.11% | 2.88% |
| Thoracic Spine | 17.53% | 15.39% | 16.52% | 6.69% |
| Lumbar Spine | 22.24% | 25.87% | 23.96% | 8.11% |
| Sacrum | 5.73% | 6.08% | 5.90% | 1.78% |
| Coccygeal Spine | 0.55% | 0.44% | 0.50% | 1.95% |
| **Thorax** | **10.15%** | **8.72%** | **9.48%** | **9.42%** |
| Clavicle | 0.88% | 0.60% | 0.75% | 0.77% |
| Ribs | 6.47% | 5.25% | 5.89% | 7.87% |
| Sternum | 2.81% | 2.87% | 2.84% | 0.79% |
| **Upper Extremities** | **11.24%** | **10.39%** | **10.84%** | **21.78%** |
| Scapula | 4.09% | 3.67% | 3.89% | 5.55% |
| Humerus | 6.89% | 6.54% | 6.72% | 6.71% |
| Forearm | 0.21% | 0.13% | 0.17% | 6.00% |
| Hand | 0.06% | 0.05% | 0.05% | 3.51% |
| **Pelvis** | **12.73%** | **12.29%** | **12.52%** | **8.02%** |
| **Lower Extremities** | **7.74%** | **9.14%** | **8.41%** | **20.53%** |
| Femur | 6.64% | 7.94% | 7.26% | 8.40% |
| Leg | 0.98% | 1.14% | 1.06% | 6.50% |
| Foot | 0.12% | 0.06% | 0.09% | 5.63% |

